# BakRep – A searchable large-scale web repository for bacterial genomes, characterizations and metadata

**DOI:** 10.1101/2024.05.28.596065

**Authors:** Linda Fenske, Lukas Jelonek, Alexander Goesmann, Oliver Schwengers

## Abstract

Bacteria are fascinating research objects in many disciplines for countless reasons, and whole-genome sequencing has become the paramount methodology to advance our microbiological understanding. Meanwhile, access to cost-effective sequencing platforms has accelerated bacterial whole-genome sequencing to unprecedented levels introducing new challenges in terms of data accessibility, computational demands, heterogeneity of analysis workflows, and thus, ultimately its scientific usability. To that end, *Blackwell et al*. released a uniformly processed set of 661,405 bacterial genome assemblies obtained from the European Nucleotide Archive as of November 2018. Building on these accomplishments, we conducted further genome-based analyses like taxonomic classification, MLST subtyping and annotation of all genomes. Here we present BakRep, a searchable large-scale web repository of these genomes enriched with consistent genome characterizations and original metadata. The platform provides a flexible search engine combining taxonomic, genomic and metadata information, as well as interactive elements to visualize genomic features. Furthermore, all results can be downloaded for offline analyses via an accompanying command line tool. The web repository is accessible via https://bakrep.computational.bio.

## Introduction

Bacteria represent a significant portion of Earth’s biodiversity, showcasing an astounding variety of habitats. For the past three decades, bacterial whole-genome sequencing (WGS) has provided deep insights into the vast diversity of populations and ecosystems’ complexity, just as into the organization and plasticity of single genomes - both fundamental for our perception of microbial life. In particular, WGS of bacterial pathogens has tremendously propelled our understanding of drug resistances, virulence factors, and host interactions and has become invaluable for medical microbiology. But simultaneously, the exploration and analysis of less-studied species continuously expands our knowledge of the broad and hard-to-comprehend diversity within the bacterial domain of life. However, the rapid and accelerating generation of WGS data demands substantial storage and analysis capacities. To securely store the raw DNA sequencing data, public databases like the Sequence Read Archive (SRA), the DNA Data Bank of Japan (DDBJ), or the European Nucleotide Archive (ENA) are primarily considered [1]. Consequently, these data repositories are in a constant state of growth. For example, at the time of writing, more than 4.6 billion sequences are stored in the ENA (https://www.ebi.ac.uk/ena/browser/about/statistics), and the latest GenBank release (v257.0) contains 2.6 billion WGS records (https://ncbiinsights.ncbi.nlm.nih.gov/2023/08/21/genbank-release-257/). Along with these rapidly growing data collections, several challenges arise with regard to the **FAIR** principles [2]. **Findability**: to conduct comparative analyses targeting particular sublineages or MLST types, sequenced samples often need to be processed prior to genome-based screening and filtering steps. **Accessibility**: the sheer amount of raw data needs to be handled and properly processed for analysis which poses a serious barrier for many researchers lacking necessary IT infrastructure and bioinformatics skills. **Interoperability**: common data formats, vocabularies and ontologies are crucial to facilitate data integration across different platforms. **Reproducibility**: large parts of this data are processed over and over again introducing adverse variability regarding used analysis tools, parameters and databases. Furthermore, user-provided metadata may be prone to inaccuracies and incompleteness complicating reproducibility and subsequent processing [3], [4]. In conclusion, this situation leads to inflated bioinformatic workloads, increasing analysis costs regarding computational resources and valuable staff time. The analyses of genomes of varying quality, being assembled and annotated using different algorithms and thus ultimately putting the usability of this valuable data at stake [5], [6]. In contrast to the large raw data repositories, dedicated initiatives, *e*.*g*., EnteroBase [7] conduct consistent data processing procedures comprising targeted and streamlined genome characterizations. However, these platforms typically focus on distinct taxa and thus, are of limited general usability. An essential step addressing these challenges was made by a previous study *by Blackwell et al*. following a uniform approach to assemble and characterize all bacterial paired-end WGS datasets retrieved from the ENA as of November 2018 [8]. As a result, 661,405 consistently assembled genomes were made publicly available facilitating the broader access and utilization of this data for the research community. This study accomplished the systematic and standardized processing of this massive dataset, and thus fostered the usability of this genomic data. However, access to these genomes remains limited, since all genome Fasta files are provided as one comprehensive 751 GB single-file archive, thus posing a significant barrier in terms of findability and accessibility for further analyses. Even though assembled genomes are pre-indexed using various search algorithms, it remains challenging for users without sufficient bioinformatics knowledge or command-line skills to find and extract genomes of interest. Hence, to fully exploit the huge potential of this highly valuable dataset, researchers would benefit from a user-friendly platform providing streamlined access to this huge amount of data via flexible search capabilities integrating the various information layers, like genome characterizations, taxonomic classifications and subtypings, annotated genomic features, and last but not least metadata. Building on these uniformly assembled bacterial genomes, here, we present BakRep, a large-scale comprehensive web repository specifically addressing these challenges. All 661,405 genomes were consistently quality controlled, taxonomically classified, multilocus sequence typed, and annotated. In line with the FAIR principles, all information is findable and accessible via an interactive website providing researchers with a versatile search engine integrating genomic and taxonomic information, annotated features, and original metadata. Batch downloads of search results can be conducted via an accompanying command line tool. BakRep is publicly available at https://bakrep.computational.bio

## Methods / Implementation

### Raw data processing

We retrieved 661,405 assemblies and associated metadata published by *Blackwell et al*. from http://ftp.ebi.ac.uk/pub/databases/ENA2018-bacteria-661k/. For taxonomic classification, the GTDB-Tk (v2.2.6) classify workflow [9] based on the Genome Taxonomy Database (GTDB) release R207 [10] was used, with the ‘--mash_db’ argument set for enabling ANI screening. Contamination and completeness of the assemblies were estimated with CheckM2 (v1.0.1) [11]. Basic statistics of the raw assemblies were collected using assembly-scan (https://github.com/rpetit3/assembly-scan). Determination of multilocus sequence types (MLST) was conducted using mlst (v2.23.0) (https://github.com/tseemann/mlst) utilizing the PubMLST database. Furthermore, assemblies were annotated with Bakta (v1.7.0) using the ‘full’ database version to use all features and the ‘-- keep-contig-headers’ flag to preserve the original contig headers of the raw assemblies [12]. Results were stored as JSON files via custom Python scripts. All analyses were implemented as part of a Nextflow [13] workflow executed in the de.NBI consortiums’ cloud computing infrastructure (https://github.com/ag-computational-bio/bakrep). The metacoder package (v0.3.6) was used for graphic summaries of the taxonomic abundances [14].

### Implementation of the web repository

The BakRep web repository is implemented as an HTTP-based API, based on Vert.x, offering public endpoints for search and data access [15]. Elasticsearch is utilized for implementing the search functionality [16]. All genomic data is stored as compressed plain text files in a S3-compatible storage. We provide a publicly available website that retrieves data via the API and visualizes it. The website’s graphical user interface is implemented as a single-page application in Vue.js 3 (https://vuejs.org). The services are deployed on a scalable Kubernetes cluster, which is currently hosted and run within the cloud computing infrastructure of the de.NBI consortium. An additional command line tool for automated large-scale downloads was implemented in Python (https://github.com/ag-computational-bio/bakrep-cli).

## Results

### Expansion of consistent genome analyses

In this study, we built up on the 661,405 bacterial genome assemblies provided by *Blackwell et al*. who uniformly assembled WGS raw data retrieved from the ENA archive as a November 2018 snapshot. We aimed to expand the range of consistent per-genome characterizations and to provide these results as accessible and userfriendly as possible. In this regard, all 661,405 assembled genomes (751 GB in total) were quality-checked and basic assembly statistics were calculated. We then taxonomically classified all genomes using the robust GTDB taxonomy, and where applicable, we further sequence-typed all genomes for which a species-specific multilocus-sequence typing schema existed. Last but not least, we performed a robust annotation of all genomes using Bakta taking advantage of its taxonomically untargeted full database version. From these 661,405 input assemblies, 648,567 were successfully characterized. To streamline the technical accessibility of all results, output files of all analysis tools were parsed, normalized, and serialized in JSON format, generating a total of 3,891,402 unique files. In addition, annotation results are also available in GenBank format, as well as nucleotide and amino acid Fasta files for all annotated coding sequences. A total of 6.15 TB of genomic information was generated and stored in a cloud-based S3 storage which is publicly available via an interactive web repository at https://bakrep.computational.bio.

### Diversity and bias across the various taxonomic ranks

Given the vast size of public databases, they naturally encompass a variety of species. Nevertheless, certain species receive more frequent attention due to their clinical relevance, ease of cultivation, or long-standing usage as model organisms. This inherent bias contributes to taxonomic imbalances in such data repositories. *Blackwell et al*. comprehensively demonstrated an intrinsic taxonomic bias at both the genus and species level [8]. However, we would like to address one more aspect: to what extent is there either bias or diversity at higher taxonomic ranks? Therefore, we comprehensively explored the distribution across all taxonomic ranks, utilizing a robust, purely genome-based, and thus objective taxonomic classification. We used GTDB-Tk, a widely utilized tool in the community, that delineates prokaryotic taxa based on systematic criteria and phylogenetic relationships using domain-specific marker genes in combination with mutual ANI-based genome distances. At the species level, and in line with former results, our analysis revealed that the 24 most prevalent species constitute 90 % of all genomes. The most abundant species were: *Salmonella enterica* (27.10 %), *Escherichia coli* (13.52 %), *Streptococcus pneumoniae* (7.80 %), *Mycobacterium tuberculosis* (7.43 %), and *Staphylococcus aureus* (7.28 %). At the genus level, the most prevalent genera were: *Salmonella* (27.99 %), *Escherichia* (13.82 %), *Streptococcus* (12.89 %), *Mycobacterium* (8.6 %), and *Staphylococcus* (7.92 %). However, despite these over-represented species and genera, the genomes contained in this repository exhibit a notable degree of diversity at higher taxonomic ranks, comprising 66 distinct phyla divided into 132 classes, 345 orders, 722 families, 2,466 genera, and 8,207 species. In comparison, the genome sequence-based GTDB database counts 175 phyla, divided into 538 classes, 1,840 orders, 4,870 families, 23,112 genera, and 107,235 species (https://gtdb.ecogenomic.org/) and the literature-based Bacterial Diversity Metadatabase (BacDive) lists 42 phyla divided into 106 classes, 255 orders, 648 families, 3,801 genera and 21,203 species (https://bacdive.dsmz.de/dashboard). Thus, this repository covers 37 % and 157 % of phyla, 24 % and 124 % of classes, 18 % and 135 % of orders, 14 % and 111 % of families, 10 % and 64 % of genera and 7 % and 38 % of species available in the genome-based GTDB and described in the literature-based BacDive databases, respectively. To illustrate both the diversity and bias of this repository, a taxonomic tree weighted by aggregated genome counts along all ranks was created (Fig. 1). For better visualization, taxa were clipped and aggregated at the family level. A more detailed version including all ranks is available in the supplemental data (Suppl. Fig 1). Notably, 1,634 assemblies (0.25 %) could not be assigned to any species epithet, of which 122 (0.02 %) could not be assigned to a genus. A closer examination of the unclassified genomes revealed that those lacking a genus assignment exhibit an average estimated completeness of only 46.50 %. Genomes lacking a species epithet classification exhibited a higher average completeness of 67.02 %, albeit with increased variability (Suppl. Fig 2).

**Figure 1:**
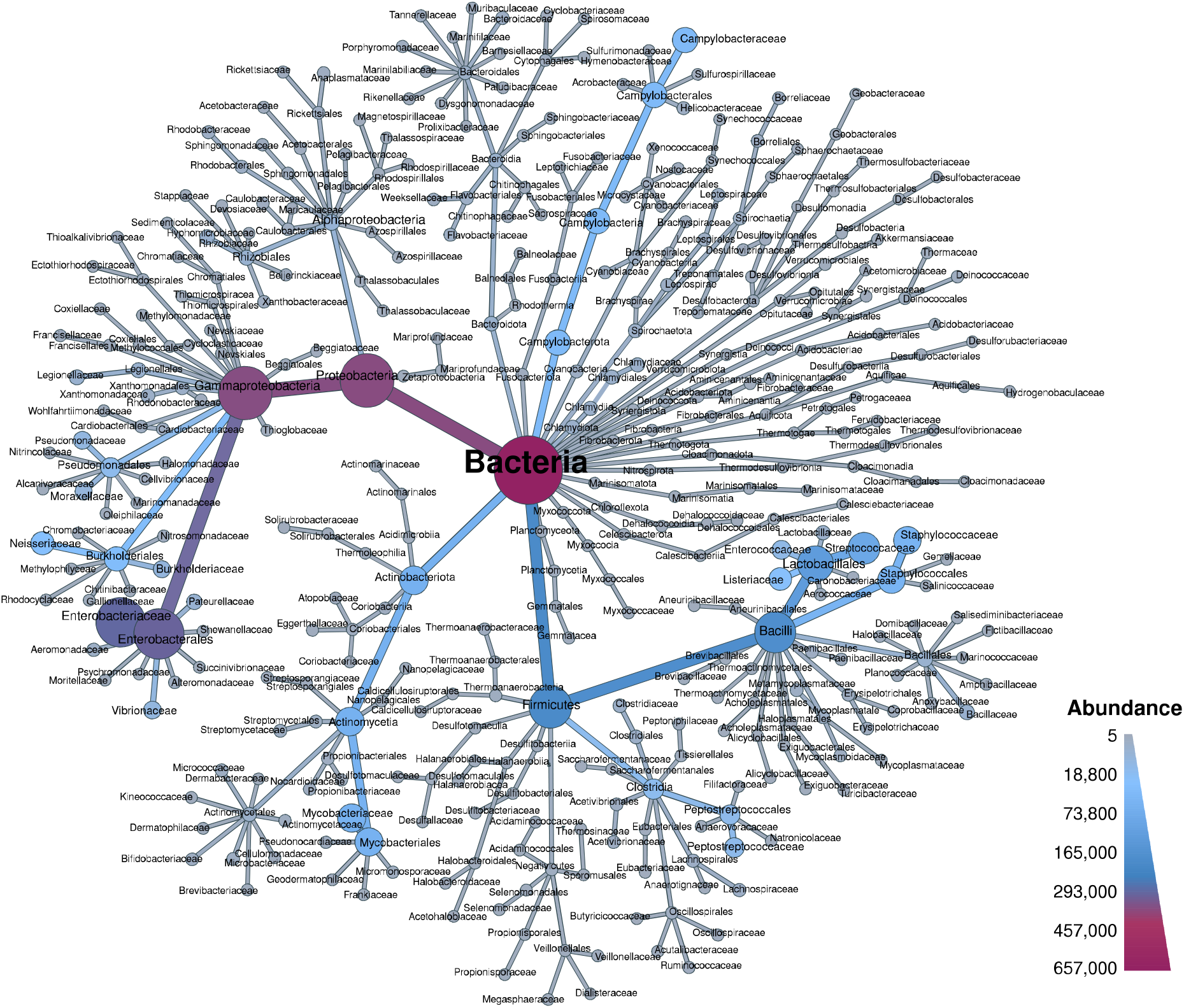
Overview of the taxonomic composition at the family level. Nodes and branches are colored and sized by aggregated genome counts at each taxonomic rank. The figure was created using the Metacoder package.

**Figure 2:**
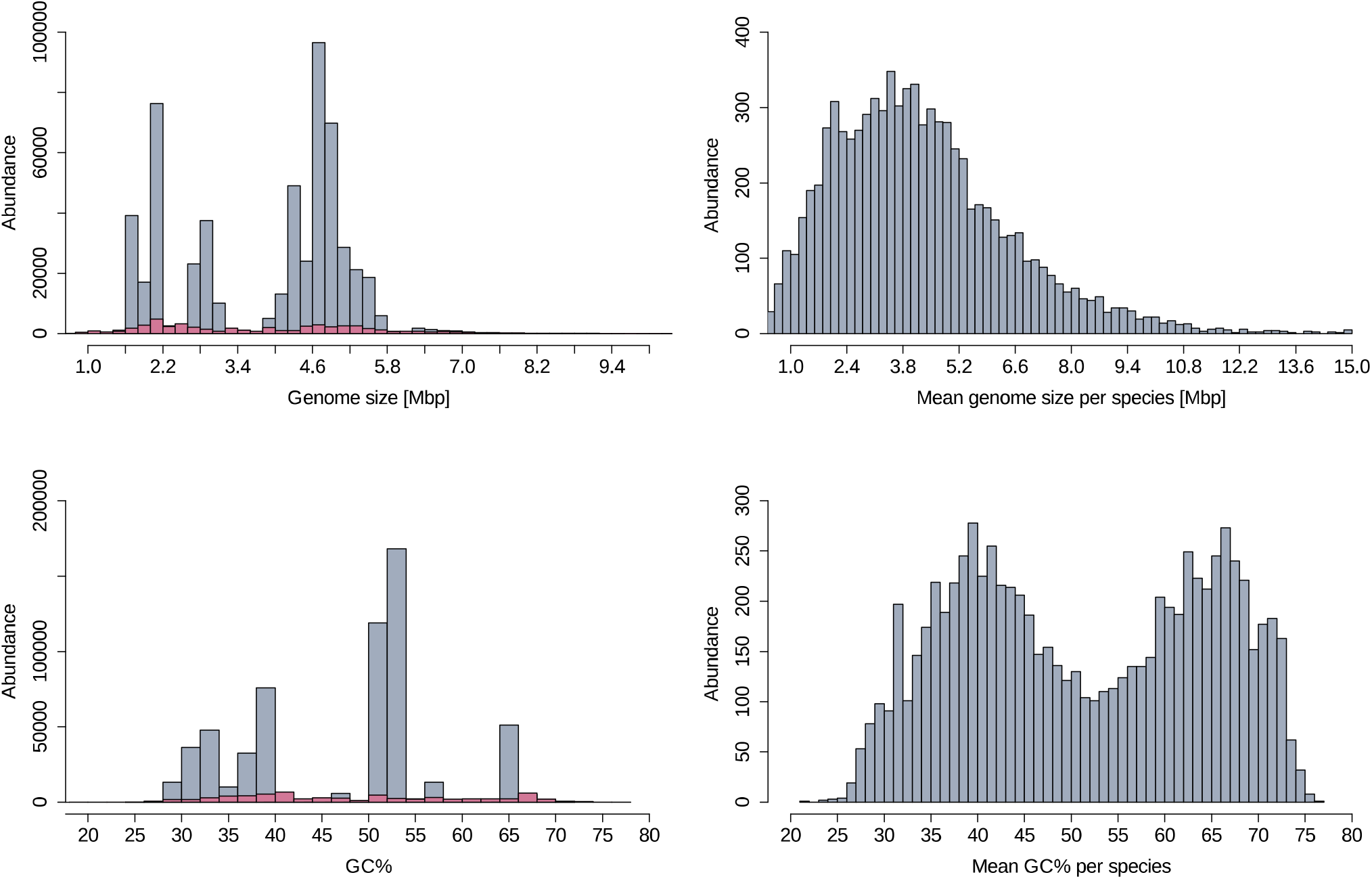
Distribution of genomic metrics in the repository. The genome size (top left) and the GC content (bottom left) are displayed for all genomes. In comparison, the mean genome size (top right) and the mean GC content (bottom right) per species are shown. The magenta-highlighted region illustrates the distribution excluding the 24 most abundant species.

Various comparative analyses begin with the selection of suitable genomes from public repositories. Here, the reliability of pre-assigned taxa is of utmost importance, immediately impacting the outcome of comparative studies. Unfortunately, user-provided taxonomic information stored as metadata in raw sequencing data archives is known to be error-prone and often does not correspond to genome-based taxonomic classifications. To address this issue, we compared the scientific taxonomic names associated with the raw data in the ENA with the genome-based taxonomic classifications conducted with GTDB-Tk. For 45,275 (6.98 %) genomes, we observed discrepancies at the species epithet. Variations at the genus level occurred in 25,913 (4.0 %) genomes. In 21,349 (3.29 %) cases, both the genus and species epithets differed. On further review, a substantial portion of these discrepancies (54.96 %) is attributed to the genus *Shigella*, which was consistently classified as *Escherichia*. Frequent inconsistencies were also evident for *Mycobacteroides abscessus*, designated as *Mycobacterium abscessus* in 2,675 cases (10.32 %), and *Burkholderia pseudomallei* classified as *Burkholderia mallei* in 1,763 cases (6.80 %). Among the 2,774 (10.71 %) species discrepancies within the *Salmonella* genus, variations arose from assigning distinct subspecies, designated as full species names by GTDB-Tk. Considering these examples, 7,106 (33.28 %) cases remained for which neither the genus nor the species epithets matched (Suppl. Tab. 1).

### Distribution of genome-based key metrics

In the NGS era, a multitude of sequencing platforms, as well as constantly evolving bioinformatics methods and implementations, contribute to a variety of assembly approaches. To quickly assess biological key features and the technical quality of assembled genomes, several metrics have evolved as gold standard indicators. For instance, the mere size of a genome alone can provide important information, for instance regarding its completeness. Also, the GC content is widely used as a rough proxy for the nucleotide composition of a genome, that is typically found in a narrow range specific to a particular bacterial species. We used the available information stored in BakRep to get an overview of the distribution of some of the most important and widely used metrics for this repository. To better understand the extent of variation within the bacterial diversity, we summarized the overall distribution of genome size and GC content. The total genome size ranges from a minimum of 100,943 base pairs (bp) to a maximum of 20,285,777 bp with a mean value of 3,901,303 bp and a median of 4,379,349 bp. To account for the observed taxonomic biases, we excluded the 24 most abundant species accounting for 90 % of all genomes. For this taxonomically clipped set of genomes, the maximum and minimum genome sizes remain unchanged, while the mean genome size increased to 3,962,704 bp, and the median decreased to 3,853,294 bp. To further mitigate the influence of over-represented regions in the genome size distribution, primarily attributed to the *Enterobacteriaceae* in the range of 4.5 - 5.5 Mbp, the *Mycobacteriaceae* in the range of 1.5 - 2.0 Mbp, and the *Vibrionaceae* and *Neisseriaceae* in the range of 2.5 - 3.5 Mbp, we calculated the mean genome size per species. In contrast, this reveals a notably homogeneous distribution with a peak at approximately 3.8 Mbp, followed by a rapid decline extending to a maximum of 15 Mbp (Fig. 2). The GC content of all genomes ranges from a minimum of 23.6 % to a maximum of 76.5 % with a mean value of 47.2 % and a median of 50.7 %. Distinct peaks are observed at approximately 40 %, within the 50 - 55 % range, and at 65 %, mostly attributed to the 24 overrepresented species. The GC content was likewise normalized based on species, resulting in a bimodal distribution that peaks at 40 % and between 60 - 70 % (Fig. 2).

In addition to genome size and GC content, further metrics evolved to quickly assess the technical quality of a sequenced and assembled genome, *e*.*g*. the number of contigs and the well-known N50 metric. However, actual values for these metrics can vary widely not only between sequencing platforms and assembly approaches, but also between species due to biological factors, like the existence and abundance of sequence repeats and mobile elements. Furthermore, due to the lack of common guidelines, it is often far from obvious which actual values are acceptable for a given metric. Hence, we leveraged the robust taxonomic classifications and vast size of this repository to aid with the provision of potential guidelines for acceptable value ranges of these key metrics per species. Hence, we examined the distributions of the aforementioned key metrics for each of the most prevalent species, including many of significant medical relevance. As anticipated, we observed substantially varying value ranges for these metrics across species (Fig. 4). Additionally, the distribution ranges within individual species also showed considerable variability. For instance, for *Klebsiella pneumoniae*, we observed notable downward deviations, with some isolates exhibiting a minimum genome size ranging from 2.8 to 4.0 Mbp, while the mean is 5.5 Mbp. However, upon closer observation, most of these outliers were identified as several isolates from the same study that utilized transposon-directed insertion-site sequencing, suggesting that these samples were not whole-genome sequenced. Discrepancies also exist for *Burkholderia mallei* with likewise noticeable downward deviations for which no clear explanation could be found within the metadata. Despite some outliers, core ranges of these key metric values might help to establish guidelines for quality assessment.

**Figure 4:**
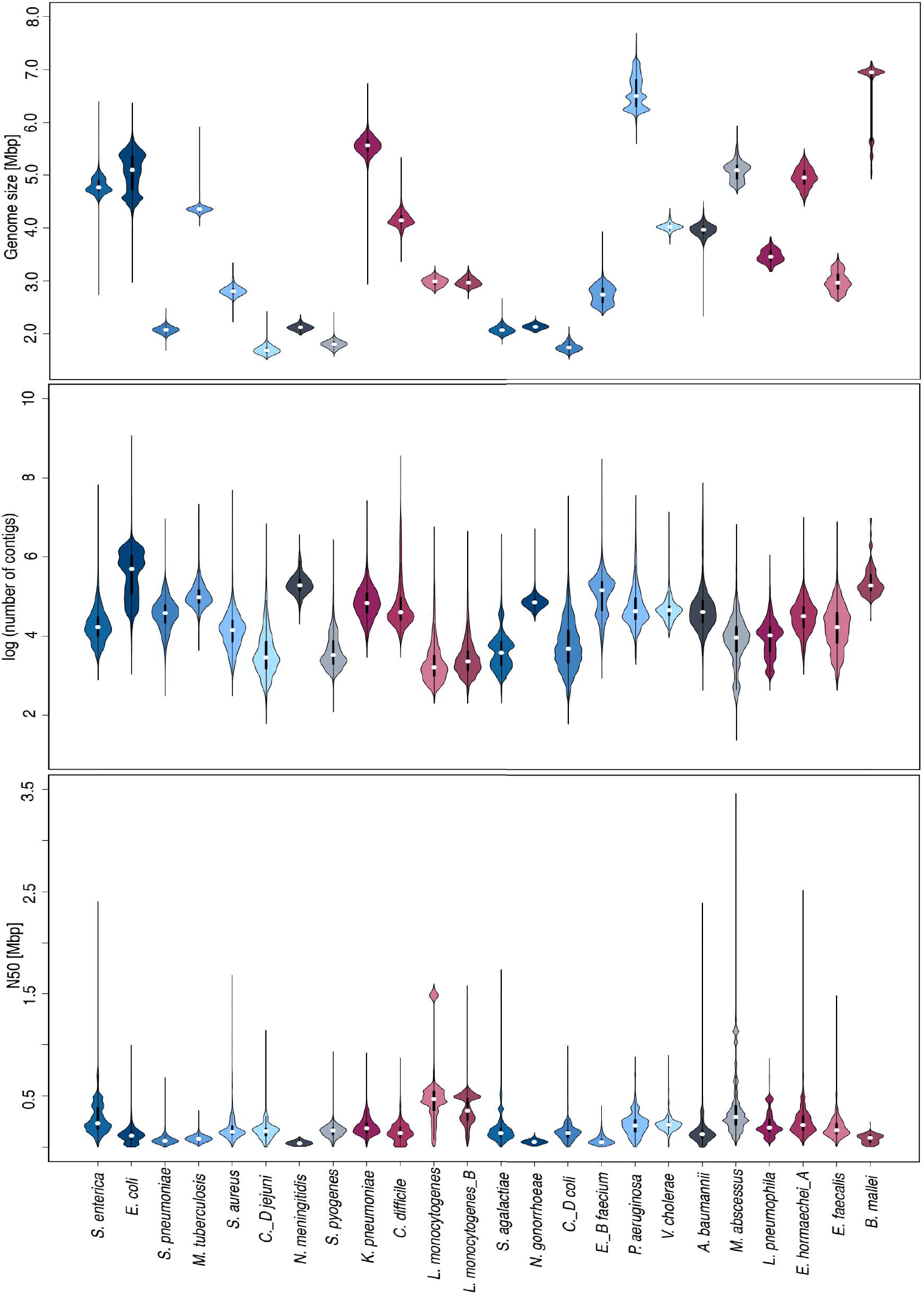
Distribution of genome assembly metrics for the 24 most abundant species. The genome size (top), the number of contigs (middle), and the N50 values (bottom) are displayed. White points indicate medians; bold black bars represent interquartile ranges; thin black lines represent outliers. Genomes were filtered for 95 % completeness and less than 1 % contamination.

The demonstrated varying ranges in genome sizes in the preceding section is an outcome of different habitats and evolutionary mechanisms constantly introducing and removing genes. Due to the intricate and diverse set of ecosystems, bacterial genomes exhibit significant variability in size and complexity, encompassing a fluid continuum between compact genomes and those with larger and more elaborate structures. As a rule of thumb, it is accepted as common knowledge that bacterial gene lengths average approximately 1 kbp per gene. To assess this assumption, we juxtaposed the mean genome sizes with the mean number of genes per species. A regression analysis revealed a slope of approximately 915 genes per 1 Mbp, resulting in a mean gene length of 1,093 bp, roughly validating but specifying this assumption with a deviation of 9.3 % and a determination coefficient (R^2^) of 0.98 confirming the postulation of a linear relationship between genome size and the number of coding genes. Besides, non-coding RNA features also play pivotal roles in bacterial genomes and cellular processes, like for example, non-coding RNAs (ncRNAs), recognized for their regulatory functions, as well as transfer and ribosomal RNAs (tRNAs/rRNAs) as essential components of the protein synthesis machinery. Hence, we likewise compared numbers of annotated non-coding RNA features to the mean genome size per species. Here, tRNAs showed a linear correlation however with significant variability (R^2^=0.56). In contrast, for ncRNAs (R^2^=0.35), and rRNAs (R^2^=0.15), no clear linear trend could be observed (Suppl. Fig. 3).

### Interactive and command line access via a searchable web repository

A major part of research would be constrained or rendered infeasible without accessible data. While a high level of standardization is crucial, it is equally essential to consider the ease of data findability and accessibility. For example, in outbreak analyses, for which the presence of specific antibiotic resistance genes is pivotal, it is essential to systematically search for genomes of a particular species characterized by distinct features such as multilocus sequence-type or virulence factors. To ensure the accessibility of our results, all data was stored in a public S3 bucket. To furthermore ensure the findability of genomes of interest, we developed and provide an interactive web page that is publicly available at https://bakrep.computational.bio. It offers diverse search and filter options, allowing and streamlining the compilation of customized cohorts. To obtain an initial comprehensive overview, all available genomes can be browsed by GC content, number of contigs, genome size, as well as estimated completeness and contamination levels. To conduct comprehensive and detailed large-scale searches, BakRep offers an advanced search engine that enables robust scalable queries flexibly combining various information like genome size, GC content, number of contigs, sequence type, different taxonomic ranks and annotated gene symbols or protein product descriptions. Furthermore, and in addition to genome analysis-based information, users also have access to quality-controlled metadata associated with each dataset upon initial raw data submission to the ENA. So, the repository supports the filtering of genomes based on various metadata, including isolation source and time, associated host species, and projekt affiliation, enabling targeted searches by criteria such as country of origin, isolation period, or host organism. A more detailed list including all possible search tags is available in the supplemental data (Suppl. Tab. 2). To name an example, in one of our ongoing research projects, we utilized this search engine to identify all *Streptococcus agalactiae* genomes that met specific quality criteria, were isolated from humans, belonged to sequence type 17, and contained the penicillin-binding proteins *pbp1a, pbp1b, pbp2a* or *pbp2X* (Fig. 5). A summary of the particular search results can be exported in TSV format. All individual genomes are displayed in human-readable formats such as a summary table, a feature table, and an igv.js-based genome browser [17], and provide cross-links to databases such as the GTDB, RefSeq [18] or UniProt [19]. Each analysis result can be accessed and downloaded per genome via the website. To facilitate extensive analyses with the download of larger genome cohorts, we offer access to the download backend through a dedicated command line tool accessible via https://github.com/ag-computational-bio/bakrep-cli.

**Figure 5:**
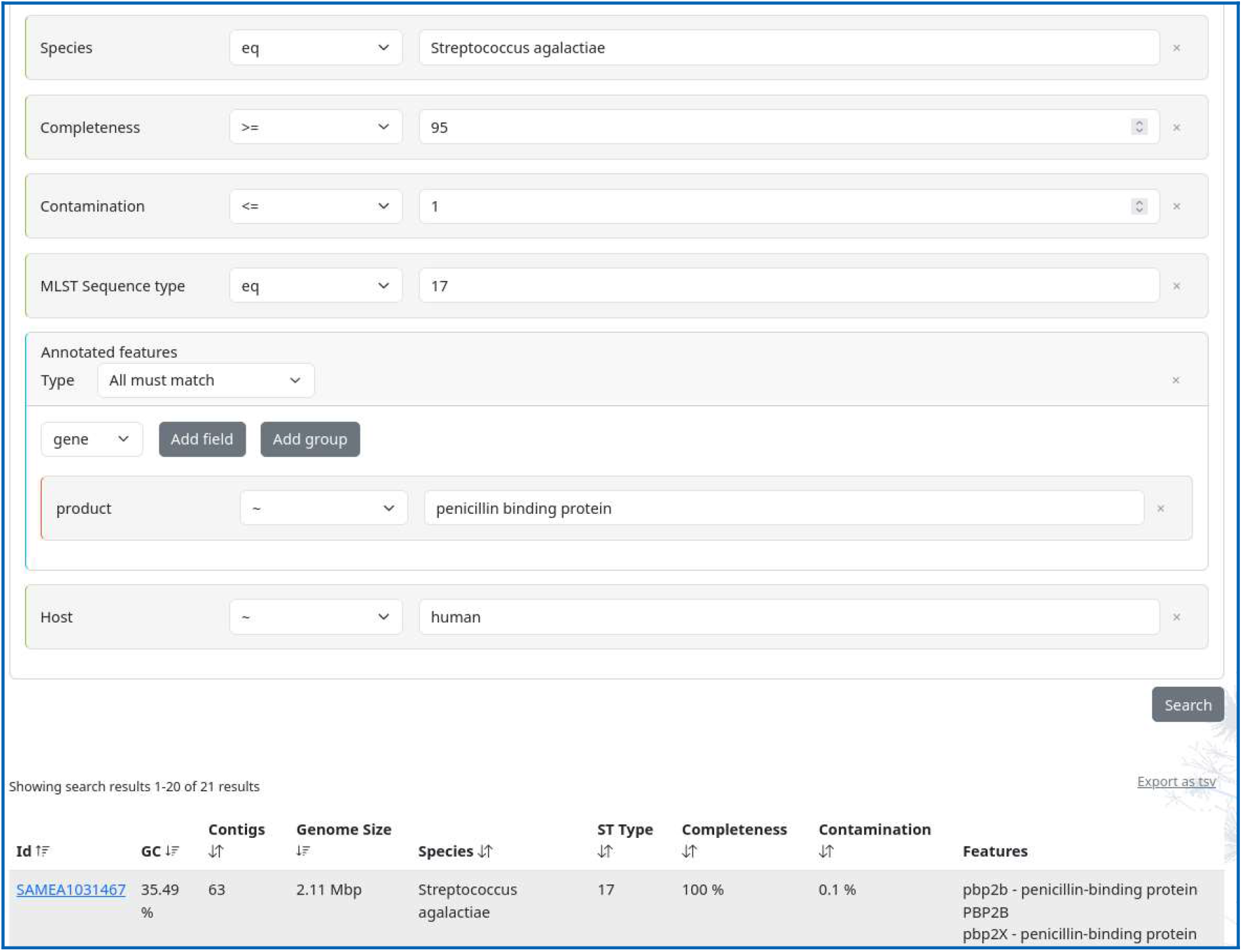
Overview of the search function of the BakRep web repository. Advanced queries for genomes with specific characteristics such as species, completeness, contamination levels, sequence type, annotated features, or host species are possible. The search outcomes are presented in a concise summary table. Details for each dataset are provided on a separate page.

## Discussion

Given the current capabilities for rapid and cost-efficient sequencing approaches, vast amounts of bacterial sequence data are generated daily. Submitting genomic data to public repositories has become a pivotal and obligatory procedure to underpin research findings and guarantee unrestricted and open access to this valuable data. Such databases consequently contain a wealth of bacterial WGS data, representing genetic reservoirs with vast potential for various applications. Nevertheless, publicly accessible genomic data often exhibit inconsistencies or insufficient processing, hindering accessibility for researchers. Challenges arise from diverse assembly methods and variations in quality control, potentially introducing batch artifacts in large-scale analyses. A critical stride in tackling these challenges was taken by *Blackwell et al*., through the uniform processing of all WGS data found in the ENA as of November 2018, which yielded 661,405 standardized assemblies. BakRep follows up on this endeavor by adding results of additional analyses, like assembly metrics, taxonomic classifications, MLST subtyping, genome annotations, and metadata of original submissions. Finally, it facilitates streamlined access to this broad amount of information via an interactive website providing powerful search capabilities.

The sequencing of bacterial genomes has become routine, significantly reshaping our understanding of the bacterial world with information gleaned from tens of thousands of genomes. Nevertheless, this quantity exhibits a notable skew toward specific phyla housing for example particular model organisms [20]. This taxonomic bias of just a few species making up the majority of genomic data may be due to various factors, such as the over-representation of well-researched and easily cultivable species, leading to gaps in the representation of the lesser researched or uncultivable microbial diversity. Furthermore, many sequencing projects focus on certain pathogens or organisms with global significance. For example, the GenomeTrakr network represents the inaugural distributed collaboration of laboratories employing WGS for pathogen identification [21], or the “10,000 Salmonella genomes project”, which sequenced more than 10,000 *Salmonella* isolates [22]. This shows the impact of funding and scientific emphasis on the diversity of sequences. In contrast to the approach employed by *Blackwell et al*., our study presents a more intricate portrayal of the taxonomic distribution through systematic species assignment utilizing the GTDB. They already acknowledged that certain aspects of sequence diversity within the assemblies might have been overlooked due to constraints inherent in the Kraken 2 database, which they used for taxonomic assignment and abundance estimation [3]. In contrast, the taxonomic classification in this study was conducted using the GTDB, employing a normalized genome-based classification derived from phylogenetic trees. These trees were constructed using a concatenated protein phylogeny, serving as the foundation for bacterial taxonomy. This approach conservatively eliminates polyphyletic groups and normalizes taxonomic ranks based on the relative evolutionary divergence [23]. However, the GTDB currently enumerates 175 phyla, divided into 538 classes, 1,840 orders, 4,870 families, 23,112 genera, and 107,235 species. This indicates that our dataset covers 37% of those phyla, 24% of classes, 18% of orders, 14% of families, 10% of genera, and 7% of species, highlighting its limited scope and underscoring that it encompasses only a fraction of the extensive bacterial diversity. There is a need to shift emphasis from a strong focus on known pathogens in sequencing projects, towards underrepresented and unknown species. This approach is crucial for a more comprehensive understanding of the patterns within bacterial diversity.

Examining several assembly metrics provides valuable insights into bacterial genomes, aiding in understanding their genetic diversity, evolutionary relationships, functional roles, and taxonomic classification. The bias of overrepresented species in the repository is also evident regarding these metrics. Nevertheless, the absence of prominently discernible gaps in the distribution of the mean genome size per species instills confidence that this snapshot may nevertheless encapsulate a substantial portion of bacterial diversity. However, while bacteria can attain genome sizes of up to 16 Mbp [24], the upper ranges are only poorly represented here. A recent study mentions a connection between the distribution of genome sizes and ecosystem type or associations with hosts, and it also discusses the ongoing challenge of precisely defining the distribution of genome size beyond the confines of laboratory settings [25]. Another study postulated an indirect mechanism of natural selection whereby ancient adaptations have induced alterations in the bacterial genome, contributing to a bimodal distribution pattern of genomic GC, which we also observed here [25]. While the genome size of a species may show some variability, caution should be exercised when encountering pronounced outliers. Substantial deviations from the mean literature value may indicate potential issues with quality, possible contamination, or the sequencing of partial segments rather than the entire genome, as exemplified in *Klebsiella pneumoniae*. Knowledge gained from a comprehensive and standardized analysis of numerous bacterial genomes has the potential to contribute valuable insights, aiding in the formulation of robust guidelines specific to certain species. Empirical values derived from diverse biological samples might offer more reliable guidelines than solely relying on literature values established over the years only using a few type strains or reference genomes. By encompassing a broader array of datasets, our repository helps to generate such guidelines. This extensive collection allows for more robust analysis and comparisons. Consequently, researchers can develop more nuanced and reliable guidelines that better reflect the complexity of bacterial genomes.

Genome fragmentation is a prevalent issue associated with short-read sequencing technologies. This challenge stems from the generation of shorter DNA fragments during the sequencing process, leading to genome assemblies typically consisting of an increased number of contigs. A recent article mentioned that the quality of these genome sequences may suffice for most analyses but need to be more practical for comparative genomics [26]. Given the typical size of a bacterial genome, a genome with a high number of contigs would result in smaller contig sizes. Referring to the average gene size of 1,093 bp, which we have calculated here, smaller contigs may contain at most one complete gene, with fragmented genes may frequently appear at contig boundaries. Significant variability in the number of contigs, especially with numerous upward outliers, should therefore, be approached cautiously. Due to the fact that the underlying assemblies in this study are based solely on Illumina sequencing, to improve the dataset it would be beneficial to include other sequencing techniques.

The presence of taxonomic misclassified species in public repositories is of significant concern to researchers, as it can introduce inaccuracies into various analyses, thus impacting the reliability of the findings. Furthermore, classification errors can propagate over time as incorrectly labeled genomes are used as references to identify novel sequences. Specific errors may stem from taxonomic naming inconsistencies or the frequent reclassification of organisms prompted by new discoveries. In our study, this applies, *e*.*g*., to the discrepancies found with the genus *Shigella*, as *Shigella* species were reclassified as later heterotypic synonyms of *Escherichia coli* in the GTDB [27]. The variations in the nomenclature of *Burkholderia* species can be similarly explained, given that *Burkholderia mallei* can be characterized as a recently evolved, host-adapted clonal lineage derived from *Burkholderia pseudomallei* [28], [29]. This may also explain the observable variations in the genome size. During host adaptation, *B. mallei* experienced considerable genome reduction [28], [29]. Given that *B. mallei* and *B. pseudomallei* share over 99 % genetic homology, taxonomic transitions between them can be fluid [30]. As GTDB-Tk uses an operational average nucleotide identity-based approach relying on type strains, only a few different genes will not lead to species differentiation. Unfortunately, we were initially unaware of the extensiveness of taxonomic discrepancies in the dataset, and thus we decided to use species information associated as metadata for our genome annotation processes. This will certainly be addressed in future versions by using GTDB-Tk species classifications ensuring accurate and consistent species listings down to annotation result files.

Adherence to the FAIR Principles - Findability, Accessibility, Interoperability, and Reusability - is crucial for advancing genomic research. With our public web repository we ensure that genomes are easily findable with persistent identifiers being accessible to a wide range of users across different platforms. By providing genome annotations in common file formats such as GenBank, we foster compatibility with various bioinformatic tools for targeted downstream analyses, thus reducing technical barriers, increasing efficiency and supporting the reproducibility of research results. Furthermore, we provide streamlined access to the valuable raw assemblies of *Blackwell et al*., ensuring that these results can be used for further studies.

## Conclusion / Outlook

The BakRep web repository provides a consistent and comprehensive characterization of one of the largest collections of bacterial genomes comprising assembly metrics, robust taxonomic classifications, MLST subtypings, genome annotations, and original metadata. Its implementation and underlying cloud infrastructure facilitate scalability and allow for swift adjustments to extended analyses and expanding datasets. Our long-term plan includes the addition of more genomes to our repository, aiming for the continuous expansion of this standardized dataset. We envision BakRep as a high-quality open resource for microbial researchers worldwide.

## Funding

This work was supported by the Justus Liebig University Giessen, Germany as well as the BMBF-funded de.NBI Cloud within the German Network for Bioinformatics Infrastructure (de.NBI) (031A532B, 031A533A, 031A533B, 031A534A, 031A535A, 031A537A, 031A537B, 031A537C, 031A537D, 031A538A).

## Acknowledgments

We gratefully thank Frank Förster for setting up the cloud project and for all the technical support. We also would like to thank Sonja Diedrich for her code contributions and the support with image editing. We gratefully thank Grace Blackwell, John Lees and Zamin Iqbal for fruitful discussions, feedback and support.

## Author contributions

OS designed and supervised the study. LJ developed the website and the servers. LF conducted data analyses, interpreted the data, and wrote the manuscript. AG supervised the study and was responsible for funding. All authors critically checked and contributed to the final version of the manuscript.

## Conflicts of interest

The authors declare that they have no conflicts of interest.

## Data availability

The website can be accessed via bakrep.computational.bio. The workflow used for data analysis is available at github.com/ag-computational-bio/bakrep. Original data was retrieved via ftp.ebi.ac.uk/pub/databases/ENA2018-bacteria-661k. The accompanying command line tool is available at https://github.com/ag-computational-bio/bakrep-cli.

## Notes

### Competing Interest Statement

The authors have declared no competing interest.

## References

[1] M. Blaxter et al., ‘Reminder to deposit DNA sequences’, Science, vol. 352, no. 6287, pp. 780–780, May 2016, doi: 10.1126/science.aaf7672.

[2] M. D. Wilkinson et al., ‘The FAIR Guiding Principles for scientific data management and stewardship’, Sci. Data, vol. 3, no. 1, Art. no. 1, Mar. 2016, doi: 10.1038/sdata.2016.18.

[3] H. Bagheri, A. J. Severin, and H. Rajan, ‘Detecting and correcting misclassified sequences in the largescale public databases’, Bioinformatics, vol. 36, no. 18, pp. 4699–4705, Sep. 2020, doi: 10.1093/bioinformatics/btaa586.

[4] F. Keck, M. Couton, and F. Altermatt, ‘Navigating the seven challenges of taxonomic reference databases in metabarcoding analyses’, Mol. Ecol. Resour., vol. 23, no. 4, pp. 742–755, 2023, doi: 10.1111/1755-0998.13746.

[5] ‘Extensive Error in the Number of Genes Inferred from Draft Genome Assemblies | PLOS Computational Biology’. Accessed: May 22, 2024. [Online]. Available: https://journals.plos.org/ploscompbiol/article?id=10.1371/journal.pcbi.1003998

[6] S. L. Salzberg, ‘Next-generation genome annotation: we still struggle to get it right’, Genome Biol., vol. 20, no. 1, p. 92, May 2019, doi: 10.1186/s13059-019-1715-2.

[7] Z. Zhou et al., ‘The EnteroBase user’s guide, with case studies on Salmonella transmissions, Yersinia pestis phylogeny, and Escherichia core genomic diversity’, Genome Res., vol. 30, no. 1, pp. 138–152, Jan. 2020, doi: 10.1101/gr.251678.119.

[8] G. A. Blackwell et al., ‘Exploring bacterial diversity via a curated and searchable snapshot of archived DNA sequences’, PLoS Biol., vol. 19, no. 11, p. e3001421, Nov. 2021, doi: 10.1371/journal.pbio.3001421.

[9] P.-A. Chaumeil, A. J. Mussig, P. Hugenholtz, and D. H. Parks, ‘GTDB-Tk v2: memory friendly classification with the genome taxonomy database’, Bioinformatics, vol. 38, no. 23, pp. 5315–5316, Dec. 2022, doi: 10.1093/bioinformatics/btac672.

[10] ‘GTDB: an ongoing census of bacterial and archaeal diversity through a phylogenetically consistent, rank normalized and complete genome-based taxonomy | Nucleic Acids Research | Oxford Academic’. Accessed: Sep. 20, 2023. [Online]. Available: https://academic.oup.com/nar/article/50/D1/D785/6370255?login=true

[11] ‘CheckM2: a rapid, scalable and accurate tool for assessing microbial genome quality using machine learning | Nature Methods’. Accessed: Sep. 20, 2023. [Online]. Available: https://www.nature.com/articles/s41592-023-01940-w

[12] O. Schwengers, L. Jelonek, M. A. Dieckmann, S. Beyvers, J. Blom, and A. Goesmann, ‘Bakta: rapid and standardized annotation of bacterial genomes via alignment-free sequence identification’, Microb. Genomics, vol. 7, no. 11, Nov. 2021, doi: 10.1099/mgen.0.000685.

[13] P. Di Tommaso, M. Chatzou, E. W. Floden, P. P. Barja, E. Palumbo, and C. Notredame, ‘Nextflow enables reproducible computational workflows’, Nat. Biotechnol., vol. 35, no. 4, pp. 316–319, Apr. 2017, doi: 10.1038/nbt.3820.

[14] ‘Metacoder: An R package for visualization and manipulation of community taxonomic diversity data | PLOS Computational Biology’. Accessed: Jan. 30, 2024. [Online]. Available: https://journals.plos.org/ploscompbiol/article?id=10.1371/journal.pcbi.1005404

[15] T. Parviainen, Real-time Web Application Development using Vert.x 2.0. Packt Publishing, 2013.

[16] C. Gormley and Z. Tong, Elasticsearch: The Definitive Guide, 1st ed. O’Reilly Media, Inc., 2015.

[17] J. T. Robinson, H. Thorvaldsdottir, D. Turner, and J. P. Mesirov, ‘igv.js: an embeddable JavaScript implementation of the Integrative Genomics Viewer (IGV)’, Bioinformatics, vol. 39, no. 1, p. btac830, Jan. 2023, doi: 10.1093/bioinformatics/btac830.

[18] N. A. O’Leary et al., ‘Reference sequence (RefSeq) database at NCBI: current status, taxonomic expansion, and functional annotation’, Nucleic Acids Res., vol. 44, no. D1, pp. D733–745, Jan. 2016, doi: 10.1093/nar/gkv1189.

[19] The UniProt Consortium, ‘UniProt: the Universal Protein Knowledgebase in 2023’, Nucleic Acids Res., vol. 51, no. D1, pp. D523–D531, Jan. 2023, doi: 10.1093/nar/gkac1052.

[20] M. Land et al., ‘Insights from 20 years of bacterial genome sequencing’, Funct. Integr. Genomics, vol. 15, no. 2, pp. 141–161, Mar. 2015, doi: 10.1007/s10142-015-0433-4.

[21] R. E. Timme, M. Sanchez Leon, and M. W. Allard, ‘Utilizing the Public GenomeTrakr Database for Foodborne Pathogen Traceback’, Methods Mol. Biol. Clifton NJ, vol. 1918, pp. 201–212, 2019, doi: 10.1007/978-1-4939-9000-9_17.

[22] M. Achtman et al., ‘Genomic diversity of Salmonella enterica -The UoWUCC 10K genomes project’, Wellcome Open Res., vol. 5, p. 223, Feb. 2021, doi: 10.12688/wellcomeopenres.16291.2.

[23] D. H. Parks et al., ‘A standardized bacterial taxonomy based on genome phylogeny substantially revises the tree of life’, Nat. Biotechnol., vol. 36, no. 10, Art. no. 10, Nov. 2018, doi: 10.1038/nbt.4229.

[24] ‘Minicystis rosea gen. nov., sp. nov., a polyunsaturated fatty acid-rich and steroid-producing soil myxobacterium | Microbiology Society’. Accessed: Jan. 30, 2024. [Online]. Available: https://www.microbiologyresearch.org/content/journal/ijsem/10.1099/ijs.0.068270-0

[25] A. Rodríguez-Gijón et al., ‘A Genomic Perspective Across Earth’s Microbiomes Reveals That Genome Size in Archaea and Bacteria Is Linked to Ecosystem Type and Trophic Strategy’, Front. Microbiol., vol. 12, p. 761869, Jan. 2022, doi: 10.3389/fmicb.2021.761869.

[26] T. H. M. Smits, ‘The importance of genome sequence quality to microbial comparative genomics’, BMC Genomics, vol. 20, no. 1, p. 662, Aug. 2019, doi: 10.1186/s12864-019-6014-5.

[27] D. H. Parks, M. Chuvochina, P. R. Reeves, S. A. Beatson, and P. Hugenholtz, ‘Reclassification of Shigella species as later heterotypic synonyms of Escherichia coli in the Genome Taxonomy Database’. bioRxiv, p. 2021.09.22.461432, Sep. 22, 2021. doi: 10.1101/2021.09.22.461432.

[28] S. Appelt et al., ‘Genetic diversity and spatial distribution of Burkholderia mallei by core genome-based multilocus sequence typing analysis’, PLoS ONE, vol. 17, no. 7, 2022, doi: 10.1371/journal.pone.0270499.

[29] L. Losada et al., ‘Continuing evolution of Burkholderia mallei through genome reduction and large-scale rearrangements’, Genome Biol. Evol., vol. 2, pp. 102–116, Jan. 2010, doi: 10.1093/gbe/evq003.

[30] C. L. Hatcher, L. A. Muruato, and A. G. Torres, ‘Recent Advances in Burkholderia mallei and B. pseudomallei Research’, Curr. Trop. Med. Rep., vol. 2, no. 2, pp. 62–69, Jun. 2015, doi: 10.1007/s40475-015-0042-2.

