## Supplementary figures and images for "BakRep – A searchable large-scale web repository for bacterial genomes, characterizations and metadata"

### Supplemental_Figure1

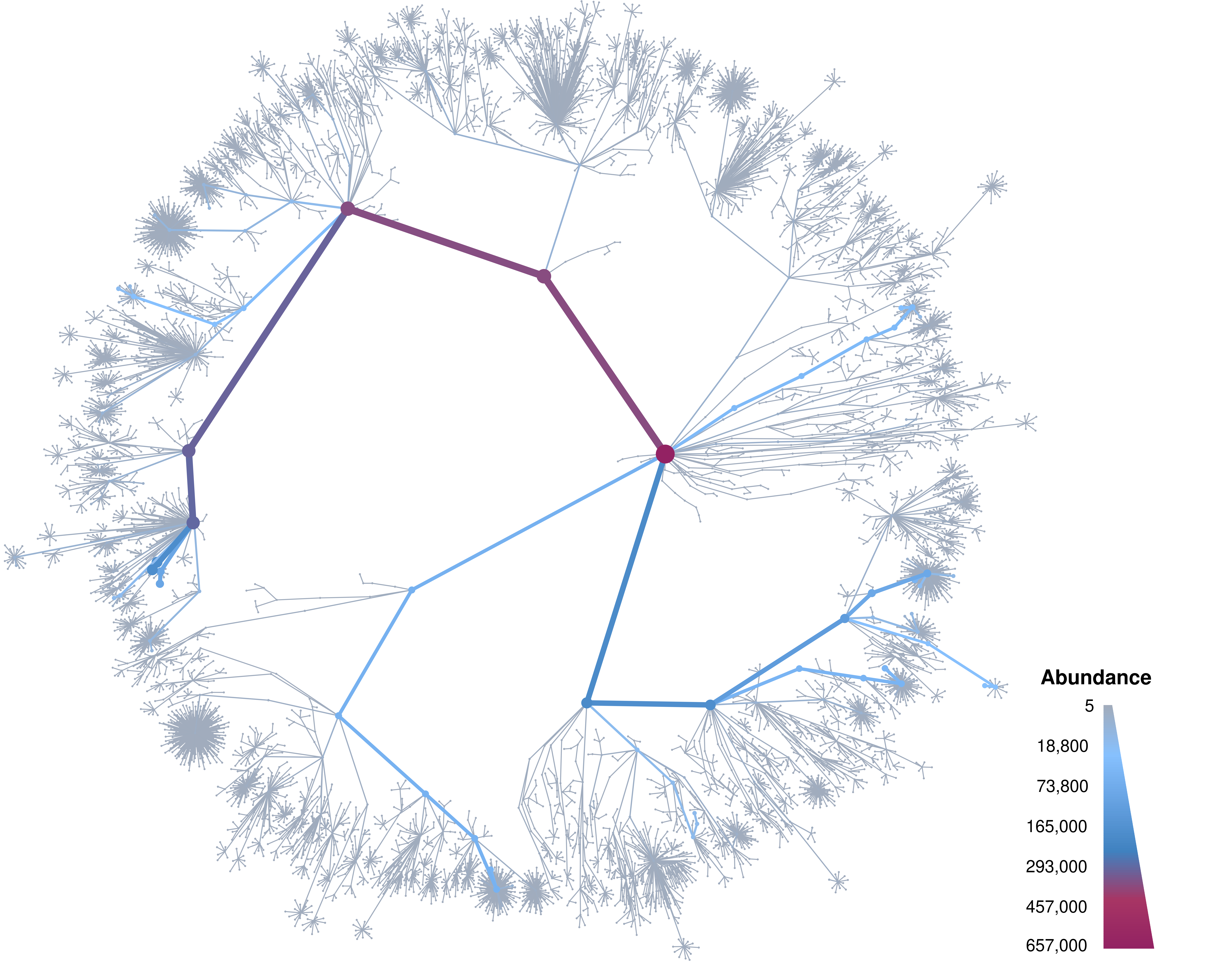

### Supplemental_Figure2

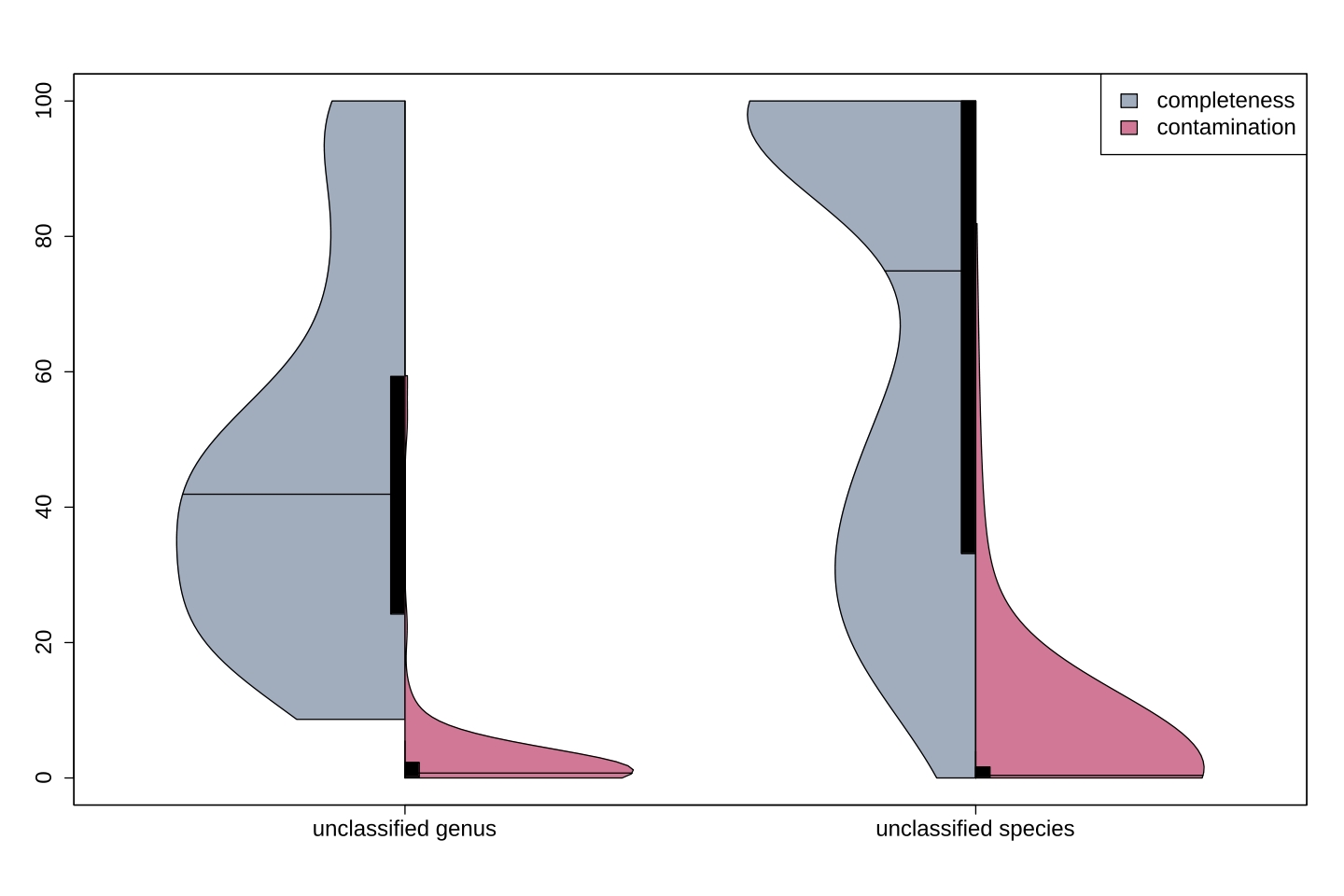

### Supplemental_Figure3

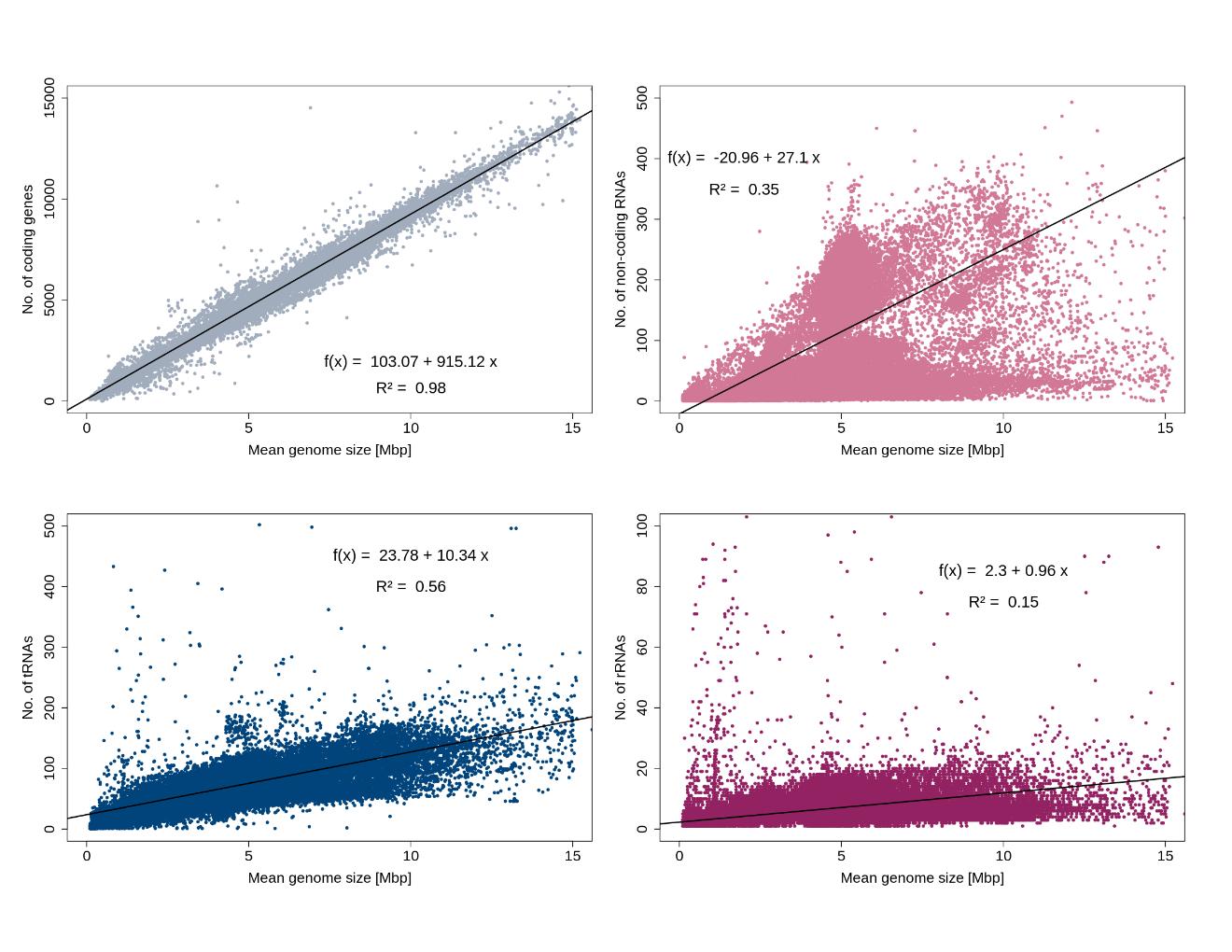
